# Development and characterization of pFluor50, a fluorogenic-based kinetic assay system for high-throughput inhibition screening and characterization of time-dependent inhibition and inhibition type for six human CYPs

**DOI:** 10.1101/2023.08.15.553391

**Authors:** Pratik Shriwas, Andre Revnew, Sarah Roo, Alex Bender, Thomas R. Lane, Sean Ekins, Craig A. McElroy

**Author notes:** **Correspondence:** Dr. Craig A. McElroy.

## Abstract

Cytochrome P450 enzymes (CYPs) play an integral role in drug and xenobiotic metabolism in humans and thus understanding CYP inhibition or activation by new therapeutic candidates is an important step in the drug development process. Ideally, CYP inhibition/activation assays should be high-throughput, use commercially available components, allow for analysis of metabolism by the majority of human CYPs, and allow for kinetic analysis of inhibition type and time-dependent inhibition. Here, we developed pFluor50, a 384-well microtiter plate-based fluorogenic kinetic enzyme assay system using substrates metabolized by six human CYPs to generate fluorescent products and determined the Michaelis-Menten kinetics constant (*K*_M_) and product formation rate (V_max_) for each substrate-CYP pair. The substrate-CYP pairs were as follows: resorufin ethyl ether for CYP1A2 (*K*_M_= 0.8 μM), CYP2C9 (*K*_M_= 0.6 μM), and CYP2D6 (*K*_M_= 2.7 μM); resorufin benzyl ether for CYP2B6 (*K*_M_= 46 μM); 3-O-methyl fluorescein for CYP2C19 (*K*_M_= 3.0 μM); and dibenzyl fluorescein for CYP3A4 (*K*_M_= 2.9 μM). We then validated each assay using known inhibitors: α-naphthoflavone for CYP1A2 (IC_50_= 13.5 nM); sertraline for CYP2B6 (IC_50_= 410 nM) and CYP2D6 (IC_50_= 2.4 μM); sulfaphenazole for CYP2C9 (IC_50_= 1 μM); ticlopidine for CYP2C19 (IC_50_= 1.2 μM); and CYP3cide for CYP3A4 (IC_50_= 56 nM). pFluor50 was also used to elucidate inhibition type and time-dependent inhibition for some inhibitors demonstrating its utility for characterizing the observed inhibition, even mechanism-based inhibition. The pFluor50 assay system developed in this study using commercially available components should be very useful for high-throughput screening and further characterization of potential therapeutic candidates for inhibition/activation with the most prevalent human CYPs.

## 2 Introduction

Cytochrome P450 enzymes (CYPs) are nearly ubiquitously expressed in numerous species across different domains (McKinnon et al., 2008) making them one of the most conserved superfamilies of proteins throughout life, present in archaea, bacteria, fungi, protists, animals and even viruses (Lamb et al., 2009). These membrane-bound proteins are expressed throughout the human body but are most abundantly expressed in the liver where they carry out biotransformations of both endogenous biomolecules such as fatty acids and steroids as well as exogenous xenobiotics (Attar et al., 2005).

The human genome comprises 57 different genes in the CYP family including genes for 18 different families and 43 subfamilies (Sadler et al., 2016, Hasler et al., 1999, Ogu and Maxa, 2000, Yan and Caldwell, 2021). However, the following six CYPs are responsible for the metabolism of more than 90% of xenobiotics in humans: CYP1A2, CYP2B6, CYP2C9, CYP2C19, CYP2D6, and CYP3A4 (Galetin et al., 2010, Zanger and Schwab, 2013, Donato et al., 2004, Miller et al., 2000, Zhao et al., 2021). Additionally, CYPs can also be responsible for drug-drug interactions in the case of co-administered drugs (Laine et al., 2001, Okudaira et al., 2007, Deodhar et al., 2020). Therefore, it is critical to understand if drug candidates are activators, inhibitors, or substrates of different CYPs (Stresser, 2004).

The drug development process is evolving due to technological innovations but is still a time-consuming and expensive process with drugs taking on average 10-12 years and 2 billion dollars of funding to get through the approval process (Hinkson et al., 2020). Additionally, there is a widely noted very high attrition rate wherein only about 10% of compounds make it through the process (Sun et al., 2022). One of the reasons for this attrition is drug metabolism and pharmacokinetics (DMPK) in which CYPs play an important role. Therefore, it is necessary to develop high-throughput methods to examine CYP-drug interactions *in vitro* early in the drug development pipeline (Ortiz de Montellano, 2013).

Currently, the most common assays for CYP inhibition are based on chromatographic techniques such as analytical HPLC, UPLC as well as LC-MS/MS analysis. These methods are laborious and time-consuming (with some assays having run times of 5-7 min/sample or more), requiring sample preparation steps prior to injection and expensive equipment as well as skilled labor (Ayrton et al., 1998, Nayadu et al., 2013). Additionally, kinetic data collection using these techniques requires a different sample for every timepoint, further decreasing the throughput. Another method that is often employed is fluorogenic or fluorescence-based assays. The main advantages of these systems are the adaptability to 384-well microtiter plates (making them high-throughput) (Sukumaran et al., 2009, Ung et al., 2018), no sample preparation required (there is no need to remove the proteins or other reagents from the reaction solution as fluorescence is emitted only by the product formed by the CYP metabolism of the substrate), and the ability to perform continuous real-time kinetic data collection. However, some drugs have intrinsic fluorescence [for example, intrinsically fluorescent chemotherapeutic agents with excitation at 350 nm and emission at 450 nm (Kabir et al., 2022)] or can cause fluorescence quenching [for example, quenching by anticancer drugs (Le et al., 2006)] so controls must be performed to ensure that the compounds being tested do not interfere with the fluorescence of the product and having non-overlapping excitation and emission wavelengths (between the fluorescent product and the drug) is essential (Paradise et al., 2007, Bell et al., 2008). Additionally, it is possible for different substrates to have distinctive inhibition or activation profiles, so having more than one fluorogenic substrate option for a given CYP can be advantageous from this perspective as well (Cohen et al., 2003).

In this study, we have developed pFluor50, a system of fluorogenic kinetic assays using substrates that form a fluorescent product through CYP metabolism with excitation wavelengths from 485-560 nM and emission wavelengths from 525-590 nM, which are non-overlapping for most intrinsically fluorescent drugs including most coumarin containing compounds (Kabir et al., 2022, Raunio et al., 2020, Ung et al., 2018). Using substrates with non-overlapping wavelengths should prevent interference from intrinsic fluorescence of drugs and also prevent quenching due to Förster resonance energy transfer. We have used the commercially available CYPexpress™ system as the source of the recombinant CYPs, which is beneficial for sample preparation if further studies of metabolite production are desired and for exploring mechanism-based inhibition as low-speed centrifugation retains the CYP system for further study or removes it for metabolite production studies. Although some of the fluorogenic substrates used have been previously reported, we report the use of resorufin ethoxy ether as a novel substrate for CYP2C9 and CYP2D6. We have determined the *K*_M_of the substrates with the corresponding CYPs and the V_max_for the formation of the product. We have further validated the assays by determining the IC_50_of known inhibitors and have demonstrated the utility of the methods for determining time-dependent inhibition and type of inhibition for some of the inhibitors.

## 3 Materials and Materials

### 3.1 Reagents

The following CYPexpress™ CYP enzyme systems [Oxford Biomedical Research, Inc.; consisting of a proprietary mix of the specific CYP as well as nicotinamide adenine dinucleotide phosphate (NADPH) reductase, magnesium (Mg^2+^), NADP+ and glucose-6-phosphate dehydrogenase (G6PDH)] were purchased from Sigma Aldrich (St. Louis, MO): CYP1A2 (Sigma-MTOXCE1A2), CYP2B6 (Sigma-MTOXCE2B6), CYP2C9 (Sigma-MTOXCE2C9), CYP2C19 (Sigma-MTOXCE2C19), CYP2D6 (Sigma-MTOXCE2D6), and CYP3A4 (Sigma-MTOXCE3A4). The following fluorogenic substrates were obtained from Cayman Chemical (Ann Arbor, MI): 7-ethoxy-3H-phenoxazin-3-one or resorufin ethoxy ether (Eres) for CYP1A2, CYP2C9, and CYP2D6 (catalog no. 16122) and 7-(benzyloxy)-3H-phenoxazin-3-one or resorufin benzyl ether (Bzres) for CYP2B6 (catalog no. 18077). The following fluorogenic substrates were purchased from Chemodex (St. Gallan, Switzerland): 2-(6-methoxy-3-oxo-3H-xanthen-9-yl)benzoic acid or 3-O-methyl fluorescein (3OMF) for CYP2C19 (catalog no.-M0098) and benzyl 2-(6-(benzyloxy)-3-oxo-3H-xanthen-9-yl)benzoate or dibenzyl fluorescein (DBF) for CYP3A4 (catalog no. – D0282). The literature precedented inhibitors were purchased from the vendors noted in parenthesis for each: α-naphthoflavone (Sigma – N5757) for CYP1A2, sertraline hydrochloride (Cayman Chemical – 14839) for CYP2B6 and CYP2D6, sulfaphenazole (Sigma – UC166) for CYP2C9, ticlopidine hydrochloride for CYP2C19 (TCI chemicals – T3110), and CYP3cide (Cayman Chemical – 15019) for CYP3A4.

Perkin Elmer (Waltham, MA – Part number 6007270) 384-well optiplate black bottom black well plates were used for all the experiments. MgCl_2_, NADPH, NADP+, and G6P were purchased from Sigma and were HPLC grade. 18.2 MΩ MilliQ water was used to prepare 0.1 M potassium phosphate buffer (pH 7.4) with potassium phosphate monobasic (0.03042 M) and potassium phosphate dibasic (0.06958 M). This buffer was used as the incubation buffer throughout the assays. The final fluorescent metabolite formation was determined using a Biotek Synergy H1 plate reader to measure fluorescence at the noted excitation (Ex) and emission (Em) wavelengths.

### 3.2 pFluor50 assay system development for CYP activity using fluorogenic substrates

The concentration of each individual CYPexpress™ reagent, the appropriate kinetic read time, and the appropriate read time interval was optimized to obtain linearity in the formation of the fluorescent product (steady-state kinetics) [Table 1 and Table S1]. Additionally, the concentration of each of the reagents was optimized for each CYP to generate an appropriate master-mix of reagents for optimal performance and linearity during the required read times. Finally, each of the plate reader parameters (Ex, Em, gain, read height, etc.) were optimized for the individual CYP-substrate pair to minimize background fluorescence and yield the highest signal-to-noise.

**Table 1.** CYP-substrate assay conditions for *K*_M_ determination.

| CYP<br>(*Amount) | Substrate | Ex:Em<br>(nm) | Read<br>time<br>(min) | Read time<br>Interval<br>(sec) | Master-mix | Substrate<br>Concentration<br>Range (μM) |
| --- | --- | --- | --- | --- | --- | --- |
| CYP1A2<br>(1 mg/ml) | Eres | 560:590 | 10 | 60 | NADPH – 0.52 mM<br>MgCl<sub>2</sub> – 2.64 mM | 0.25 – 3 |
| CYP2B6<br>(1 mg/ml) | Bzres | 535:590 | 60 | 90 | NADPH – 0.33 mM<br>MgCl<sub>2</sub> – 3.3 mM<br>NADP+ – 1.3 mM<br>G6P- 3.3 mM<br>G6PDH - 0.4 unit/ml | 1.25 – 80 |
| CYP2C9<br>(1 mg/ml) | Eres | 535:590 | 40 | 120 | NADPH – 0.65 mM<br>MgCl<sub>2</sub> – 3.3 mM | 0.25 – 8 |
| CYP2C19<br>(2 mg/ml) | 3OMF | 485:525 | 20 | 30 | NADPH – 0.33 mM<br>MgCl<sub>2</sub> – 3.3 mM<br>NADP+ – 1.3 mM<br>G6P- 3.3 mM<br>G6PDH - 0.4 unit/ml | 1 – 12 |
| CYP2D6<br>(1 mg/ml) | Eres | 560:590 | 75 | 90 | NADPH – 2.6 mM<br>MgCl<sub>2</sub> – 3.3 mM | 1 – 32 |
| CYP3A4<br>(1 mg/ml) | DBF | 485:535 | 20 | 30 | NADPH – 0.33 mM<br>MgCl<sub>2</sub> – 3.3 mM<br>NADP+ – 1.3 mM<br>G6P- 3.3 mM<br>G6PDH - 0.4 unit/ml | 0.25 – 6 |

\* - Indicated amount is the amount of CYPExpress™ reagent

In general, the protocol for determination of the Michaelis-Menten kinetic parameters (*K*_M_ and V_max_) involved addition of the appropriate master-mix to phosphate buffer followed by addition of the appropriate amount of enzyme to the mix. 30 μl of this enzyme mix was then added into each well of the 384-well plate. Next, initial background readings were taken for 5 minutes, after which the appropriate amount of substrate was dispensed into each well using the onboard reagent dispenser of the plate reader. All plates contained one blank well without enzyme (only master-mix) for each substrate concentration. Fig. S1 shows the layout of the well plate for determining the *K*_M_ for CYP3A4 as an example. Table 1 provides the formulation of the master-mix for each CYP, the kinetic read time with substrate, the read time interval, the concentration range of the substrate used for the Michaelis-Menten kinetics, the Ex and Em wavelengths, and the concentration of substrate for inhibition/activation screening. Table S1 details the specific parameters used for the plate reader for each CYP-substrate pair.

### 3.3 Michaelis-Menten kinetics for each CYP-substrate pair

For determining the Michaelis-Menten kinetic parameters (*K*_M_ and V_max_) for each CYP-substrate pair, kinetic studies were performed in which a range of different concentrations of the substrate were used (Atkins, 2005). Table 1 details the formulations for the master-mix, CYPexpress™ concentrations, and concentration ranges of each specific substrate. For each CYP-substrate pair, plate reader parameters were set up as described in Table S1. The protocol used for Michaelis-Menten kinetics was similar for all the CYPs. Briefly, the working solution was prepared for each substrate and then primed into one of the dispensers on the plate reader. The other dispenser was primed with the buffer solution. 1.5 ml of phosphate buffer solution (pH 7.4) was used to prepare the master-mix with the appropriate concentration of each reagent as specified in Table 1 (NADPH, NADP+, G6P, G6PDH, and/or MgCl_2_). The appropriate amount of each CYPexpress™ reagent was measured using an analytical balance and was transferred to the master-mix solution. This solution was mixed and 30 μl of the solution was added to each well using a multichannel pipette. In each case, a blank control well was also included as described in Fig. S1. The plate was then transferred to the plate reader and readings were taken at the time intervals, for the appropriate length of time, and at the appropriate wavelengths as specified in Table 1.

### 3.4 Determination of IC_50_values for each CYP specific inhibitor

The CYP inhibition studies used the concentrations of substrate selected based on the *K*_M_ values obtained in the kinetics studies above and are detailed in Table 2. The conditions of the inhibition studies were similar to those used in the kinetics experiments for the master-mix formulation, enzyme amount, and assay time, but with different concentrations of inhibitors added (Table 2). It is noted that the control inhibitors selected had different mechanisms of action (some have a time-dependence while others do not), so the incubation time for the enzyme-inhibitor interaction was also optimized (Table 2) and inhibition was allowed to occur for the appropriate time prior to addition of the substrate. The solvent used for each inhibitor is also noted in Table 2, as well as the final concentration of the solvent which was maintained for each concentration tested. The same amount of solvent was added to the positive and negative control to ensure that any solvent effects were appropriately normalized.

**Table 2.** CYP-substrate assay conditions for IC50 determination.

| CYP<br>(Amount) | Substrate<br>(Concentration) | Inhibitor | Solvent<br>(Final<br>Concentration) | Pre-Incubation<br>Time<br>(min) | Inhibitor<br>Concentrations<br>($\mu\text{M}$) |
| --- | --- | --- | --- | --- | --- |
| CYP1A2<br>(1 mg/ml) | Eres<br>(2 $\mu\text{M}$) | α-<br>Naphthoflavone | DMSO<br>(1%) | *10 | 0.00001 – 1 |
| CYP2B6<br>(1 mg/ml) | Bzres<br>(10 $\mu\text{M}$) | Sertraline | Methanol<br>(0.5%) | 30 | 0.0003 – 30 |
| CYP2C9<br>(1 mg/ml) | Eres<br>(2 $\mu\text{M}$) | Sulfaphenazole | DMSO<br>(1%) | *10 | 0.003 – 300 |
| CYP2C19<br>(2 mg/ml) | 3OMF<br>(2 $\mu\text{M}$) | Ticlopidine | Acetonitrile<br>(1.67%) | 5 | 0.0003 – 300 |
| CYP2D6<br>(1 mg/ml) | Eres<br>(2 $\mu\text{M}$) | Sertraline | Methanol<br>(0.5%) | 30 | 0.001 – 100 |
| CYP3A4<br>(1 mg/ml) | DBF<br>(4 $\mu\text{M}$) | CYP3cide | DMSO<br>(0.5%) | 5 | 0.001 – 100 |

\* - Time Independent

### 3.5 Determination of product formation rate

Calibration curves were constructed using the determined plate reader settings (Fig. S3) and concentrations of the products formed to allow for the calculation of product formation rates based on the observed fluorescence. For these calculations, we presumed that the total increase in observed fluorescence over time was due only to product formation. Calibration curves were generated with 6 different concentrations of the products (fluorescein and resorufin) which spanned the relative fluorescence units (RFU) observed during the kinetic experiments. This allowed for the calculation of product concentrations from the observed RFU, enabling the calculation of the product formation rates. Unfortunately, fluorescein benzyl ester (the product formed from metabolism of dibenzyl fluorescein) is not commercially available. Therefore, the exact rate of formation could not be determined and the V_max_for CYP3A4 cleavage of dibenzyl fluorescein has been reported as the rate of change in relative fluorescence units (RFU) per minute.

### 3.6 Data analysis

Slopes were calculated from the collected data by determining the increase in RFUs with respect to time. For the determination of kinetic parameters (*K*_M_ and V_max_) the slopes were then used as the input for non-linear regression fitting using a Michaelis-Menten kinetics model for a single enzyme. For determining the IC_50_of the inhibitors, the slopes were used as the input for non-linear regression fitting using the following equation (Beam and Motsinger-Reif, 2014)

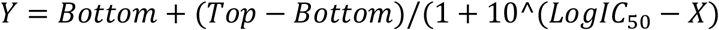

where X = the log of the substrate concentration, Y is the change in fluorescence intensity (slope), Top is the slope at full activity (uninhibited), and Bottom is the slope when fully inhibited.

All data analysis was performed using Prism 9.0 software (Graphpad Software Inc.). Experiments were performed with three biological replicates of four technical replicates each per experimental condition and the values reported are the average of these experiments. Single-factor analysis of variance (ANOVA) was used to determine statistical significance and established with P-value (\**P* ≤ 0.05, \*\**P* ≤ 0.01, and \*\*\**P* ≤ 0.001).

## 4 Results

### 4.1 Calibration curves for different substrates

The calibration curves corresponding to two of the three products (fluorescein and resorufin) along with the gain values used can be seen in Fig. S3A-F. The gain values used in the calibration curves were maintained in all other assays. All other data has been normalized to the corresponding product formation rates based on these calibration curves. Fluorescein benzyl ester (the product formed from dibenzyl fluorescein) is not commercially available so the product formation rate could not be determined and the V_max_for CYP3A4 cleavage of dibenzyl fluorescein has been reported as the rate of change in relative fluorescence units (RFU) per minute.

### 4.2 Michaelis-Menten analysis for determination of kinetic parameters of CYP-substrate pair

Michaelis-Menten kinetics analysis was performed using different concentrations of the substrates and the *K*_M_ values were determined (Table 3). The *K*_M_ values determined were comparable to those reported in the literature where available. Eres is a known substrate for CYP1A2 and the *K*_M_ value of 0.78 ± 0.3 μM is similar to the reported value of 0.62 ± 0.14 μM (Murayama et al., 2004). Bzres was used as a substrate for CYP2B6 and the *K*_M_ value of 23.6 ± 4.8 μM was close to the previously determined value of 34.0 ± 10.4 μM (Appiah-Opong et al., 2007). 3OMF is a known substrate for CYP2C19 and the *K*_M_ value was found to be 3.1 ± 0.8 μM. This value is similar to the previously reported *K*_M_ value of 1.1 ± 0.9 μM for OMF metabolism by CYP2C19 (Sudsakorn et al., 2007). DBF is a known substrate for CYP3A4 and the *K*_M_ value reported here, 1.35 ± 0 .6 μM, is comparable to the literature value of 0.87 ± 0.12 μM (Stresser et al., 2000). We demonstrate here that Eres is also a substrate for CYP2C9 with a *K*_M_ of 0.42 ± 0.1 μM. Similarly, Eres is a novel substrate for CYP2D6 with a *K*_M_ of 2.91 ± 0.8 μM. Thus, the fluorogenic method developed here uses Eres as a novel substrate for CYP2C9 and CYP2D6. Further, the V_max_was determined for the formation of the product. It was assumed that the increase in fluorescence observed was only due to the formation of the final product. As discussed above, a dose dependent fluorescence calibration curve was generated for each final product and used to calculate the concentration of product at each timepoint (with the exception of fluorescein benzyl ester). V_max_values were then calculated from the concentration of the final product at each timepoint yielding the rate of formation. Table 3 details the average V_max_for each CYP-substrate pair. Overall, the Michaelis-Menten kinetics analysis demonstrated *K*_M_ values similar to those reported in the literature for CYP1A2, CYP2B6, CYP2C19, and CYP3A4 (Appiah-Opong et al., 2007, Murayama et al., 2004, Stresser, 2004, Sudsakorn et al., 2007) whereas novel probes were identified for CYP2C9 and CYP2D6. Fig. 1 shows Michaelis Menten kinetic parameters for each CYP-substrate pair.

**Table 3.** Michaelis Menten kinetic parameters for different CYP-substrate pairs.

| CYP<br>(Substrate) | Average $V_{\max}$<br>(nmol/min) | Average $K_M$<br>($\mu\text{M}$) | Literature $K_M$<br>($\mu\text{M}$) | Reference |
| --- | --- | --- | --- | --- |
| CYP1A2<br>(Eres) | $1.83 \pm 0.4$ | $0.78 \pm 0.3$ | $0.62 \pm 0.14$ | (Murayama et al., 2004) |
| CYP2B6<br>(Bzres) | $0.62 \pm 0.21$ | $23.6 \pm 4.8$ | $34.0 \pm 10.4$ | (Appiah-Opong et al., 2007) |
| CYP2C9<br>(Eres) | $0.42 \pm 0.0$ | $0.42 \pm 0.1$ | NA | NA |
| CYP2C19<br>(3OMF) | $19.3 \pm 1.9$ | $3.1 \pm 0.8$ | $1.1 \pm 0.9$ | (Sudsakorn et al., 2007) |
| CYP2D6<br>(Eres) | $1.00 \pm 0.1$ | $2.91 \pm 0.8$ | NA | NA |
| CYP3A4<br>(DBF) | $740.2 \pm 167.9$<br>(RFU/min) | $1.35 \pm 0.6$ | $0.87 \pm 0.12$ | (Stresser et al., 2000) |

NA - Not applicable

**Figure 1.**
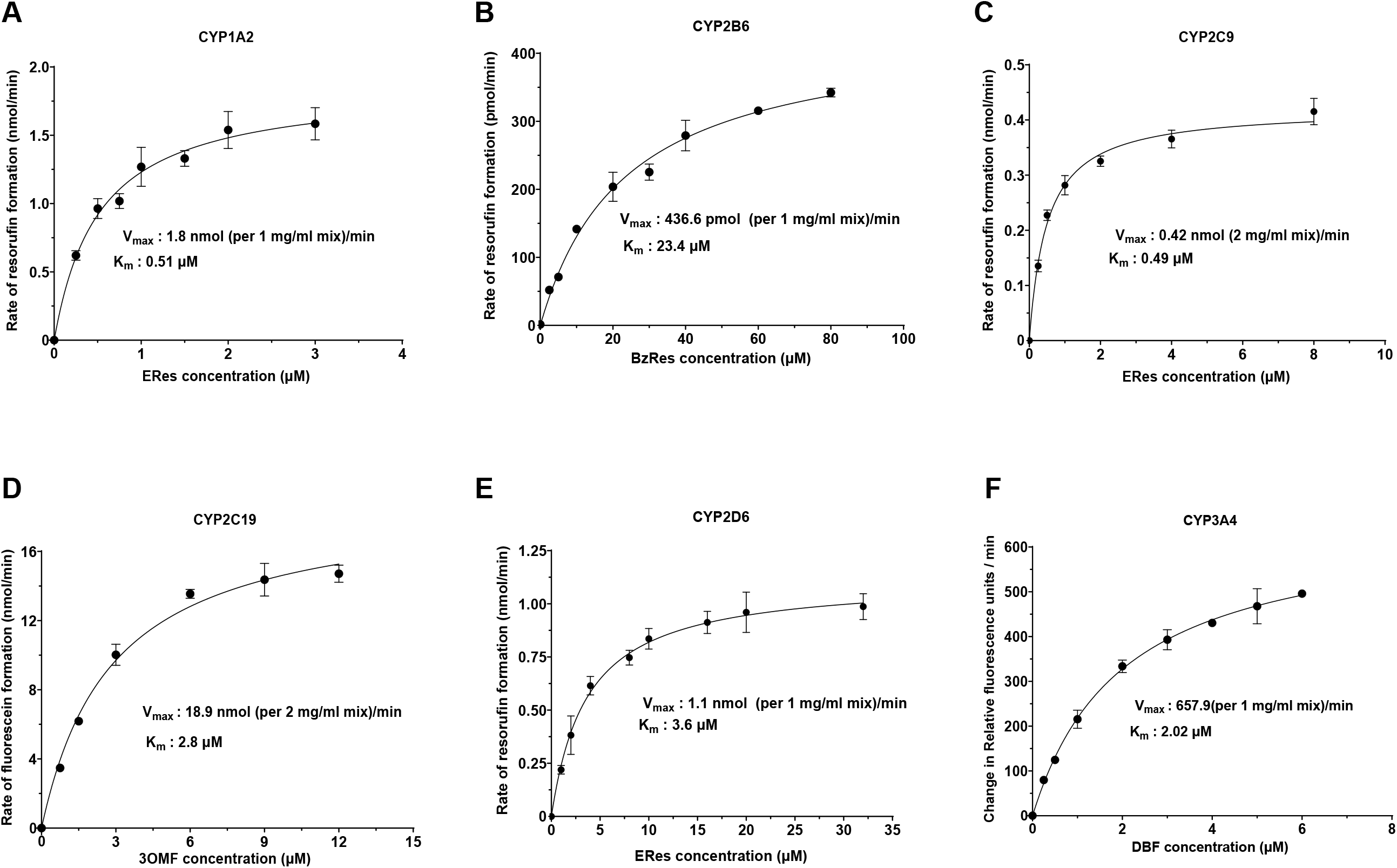
Michaelis-Menten kinetic analysis of the 6 CYP-substrate pairs. (A) Eres was used as a substrate for CYP1A2 with a *K*_M_ of 0.51 μM and a Vmax of 1.8 nmol/min (with a 1 mg/ml enzyme mix). (B) Bzres was used as a substrate for CYP2B6 with a *K*_M_ of 23.4 μM and a Vmax of 436.6 pmol/min (with a 1 mg/ml enzyme mix). (C) Eres was used as a substrate for CYP2C9 with a *K*_M_ of 0.49 μM and a Vmax of 0.42 nmol/min (with a 1 mg/ml enzyme mix). (D) 3OMF was used as a substrate for CYP2C19 with a *K*_M_ of 2.6 μM and a Vmax of 16.9 nmol/min (with a 2 mg/ml enzyme mix). (E) Eres was used as a substrate for CYP2D6 with a *K*_M_ of 3.6 μM and a Vmax of 1.1 nmol/min (with a 1 mg/ml enzyme mix). (F) DBF was used as a substrate for CYP3A4 with a *K*_M_ of 2.0 μM and a Vmax of 657 RFUs/min (with a 1 mg/ml enzyme mix). *K*_M_ and Vmax values presented in the figure are for the biological replicate shown. Each biological replicate consists of 4 technical replicates. All experiments have been performed as three biological replicates and average and standard deviation of the KM and Vmax values are presented in Table 3.

### 4.3 Determination of the time dependence of CYP inhibition by specific inhibitors

Interestingly, it was found that for specific CYPs, some of the selected inhibitors (which are FDA approved drugs) were time dependent, showing statistically significant differences in inhibition depending on the pre-incubation time whereas others were independent of time (Fig. 2). Therefore, the pre-incubation time between the inhibitor and the CYP was first optimized prior to the full IC_50_determinations. Fig. 2 details the time dependence for each of the CYPs with the selected inhibitors. Fig. 2A shows that 30 nM α-naphthoflavone inhibits CYP1A2 in a time independent manner as was noted previously (Thomford et al., 2016). Therefore, a 10-minute pre-incubation time was selected for convenience. It is documented that sertraline is a substrate for CYP2B6 and is converted to N-desmethylsertraline, which can also be further metabolized (Obach et al., 2005). Fig. 2B demonstrates that the inhibition of CYP2B6 with 30 μM sertraline increases with time, suggesting that the sertraline metabolites are likely more inhibitory towards CYP2B6. However, at short pre-incubation times with sertraline, the variability in the IC_50_data was high suggesting that the metabolism of sertraline is too fast to determine direct inhibition by sertraline without interference from the metabolites. Therefore, the sertraline pre-incubation time for the inhibition assay was optimized to 30 minutes (the time at which inhibition was no longer time-dependent). Fig. 2C demonstrates that 1 μM sulfaphenazole inhibits CYP2C9 in a time independent manner as previously established (Tseng et al., 2014) so a 10-minute pre-incubation time was selected for convenience.

**Figure 2.**
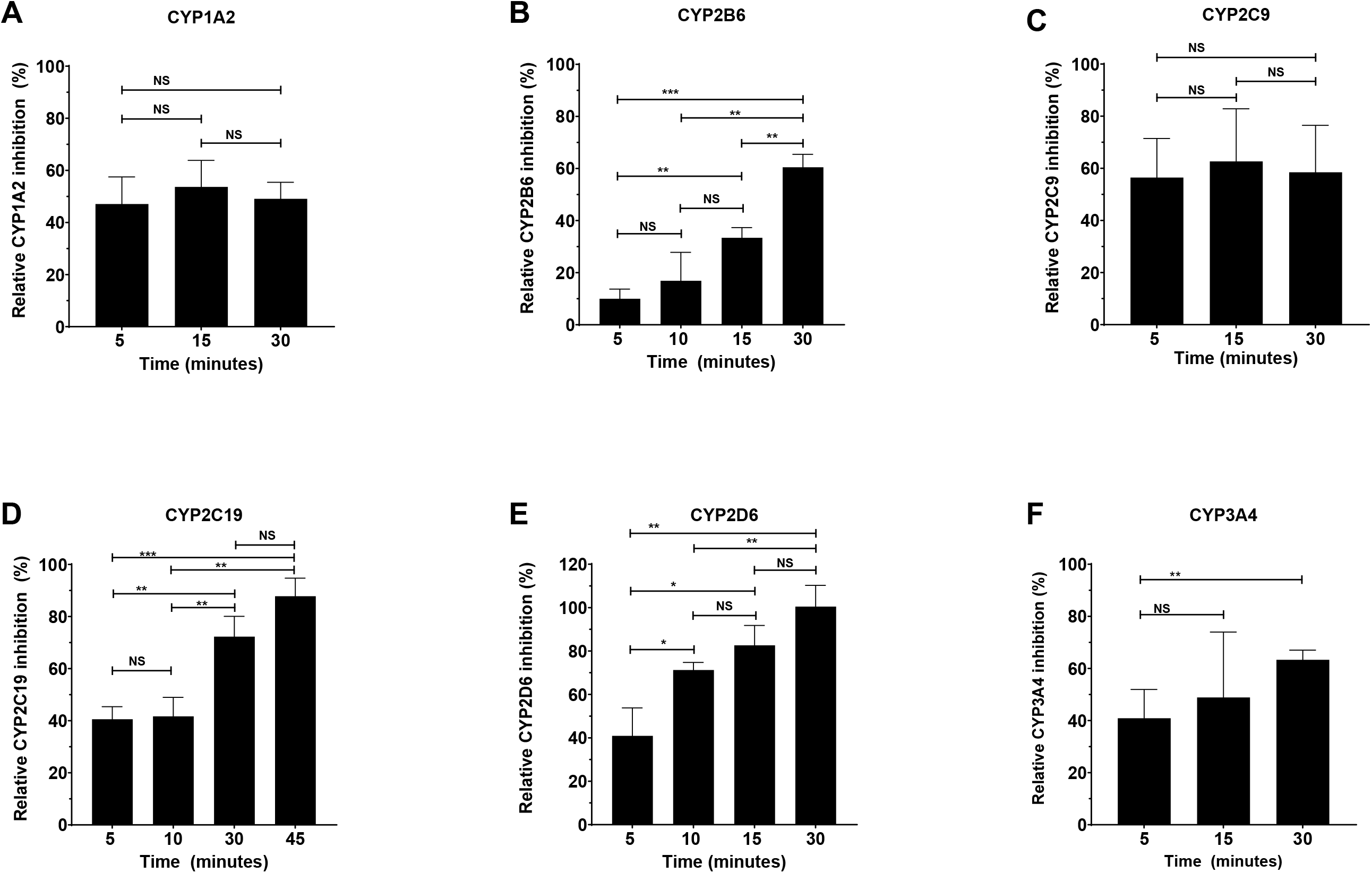
Time dependent inhibition of CYPs using specific inhibitors. (A) CYP1A2 was inhibited by 0.03 μM α-naphthoflavone in a time independent manner between 5-30 minutes. (B) CYP2B6 was inhibited by 30 μM sertraline in a time dependent manner between 5-30 minutes. (C) CYP2C9 was inhibited by 1 μM sulfaphenazole in a time independent manner between 5-30 minutes. (D) CYP2C19 was inhibited by 0.3 μM ticlopidine in a time dependent manner between 5-45 minutes. (E) CYP2D6 was inhibited by 30 μM sertraline in a time dependent manner between 5-30 minutes. (F) CYP3A4 was inhibited by 0.05 μM CYP3cide in a time dependent manner between 5-30 minutes. All the experiments have been performed as three biological replicates. Each biological replicate consists of 4 technical replicates. Single factor ANOVA was used for determination of statistical significance (NS-Not significant, *P ≤ 0.05, **P ≤ 0.01, and ***P ≤ 0.001).

Ticlopidine is known to be a time dependent inhibitor as it is metabolized to an active metabolite through the action of CYP2C19 (Nishiya et al., 2009). We found that the inhibition of CYP2C19 by 0.3 μM ticlopidine increased with increasing time (Fig. 2D). To ensure that the inhibition being measured was due to ticlopidine rather than its metabolite, 5 minutes (the shortest time attainable with reagent additions and mixing) was selected as the optimal time for the IC_50_inhibition study to determine the direct inhibition by ticlopidine. Similarly, Fig. 2E demonstrates that an increase in inhibition with time was also observed for 30 μM sertraline with CYP2D6 suggesting that CYP2D6 also metabolizes sertraline (Obach et al., 2005) and too quickly for measurement of direct inhibition. Thus, a 30-minute pre-incubation time was also selected for CYP2D6 inhibition. It is well known that CYP3cide is a mechanism-based inhibitor of CYP3A4 and Fig. 2F demonstrates that inhibition of CYP3A4 by 50 nM CYP3cide does indeed increase with time. Again, a pre-incubation time of 5 minutes was selected to determine direct inhibition by CYP3cide (Walsky et al., 2012). However, at longer incubation times, a shift in the IC_50_was observed demonstrating the utility of the method for characterizing the rate of irreversible inactivation (data not shown).

### 4.3 Determination of the IC50 for known inhibitors of each CYP

Once the pre-incubation times were optimized, full IC_50_curves were generated for each of the six CYPs with the selected inhibitors. Fig. 3 shows the IC_50_curves as well as the calculated IC_50_values for each of the inhibitor-CYP pairs and Table 4 details the average and standard deviation of the IC_50_values from three biological replicates of four technical replicates each (the data for each individual repeat is included in Table S4) and compares these values to literature data, showing good general agreement.

**Table 4.** Average IC50 values for each CYP-inhibitor pair.

| CYP<br>(Inhibitor) | Average IC<sub>50</sub><br>($\mu\text{M}$) | Literature IC<sub>50</sub> ($\mu\text{M}$)<br>(Reference)<br>[Substrate Used / Method] |
| --- | --- | --- |
| CYP1A2<br>(α-Naphthoflavone) | $0.014 \pm 0.0013$ | $0.0061 \pm 0.0008$<br>(Park et al., 2022)<br>[Phenacetin / LC-MS/MS] |
| CYP2B6 | $0.48 \pm 0.1$ | $0.20 \pm 0.024$ |
| (Sertraline) |  | (Otton et al., 1996)<br>[Bupropion / LC-MS/MS] |
| CYP2C9<br>(Sulfaphenazole) | $0.92 \pm 0.1$ | 0.6<br>(Ha-Duong et al., 2001)<br>[Teinilic acid / LC-MS/MS] |
| CYP2C19<br>(Ticlopidine) | $1.33 \pm 0.3$ | $3.7 \pm 0.7$<br>(Donahue et al., 1997)<br>[(S)-Mephenytoin / LC-MS/MS] |
| CYP2D6<br>(Sertraline) | $2.06 \pm 0.3$ | $4.35 \pm 0.5$<br>(Jin et al., 2016)<br>[Primaquine / LC-MS/MS] |
| CYP3A4<br>(CYP3cide) | $0.11 \pm 0.02$ | $0.30 \pm 0.02$<br>(Walsky et al., 2012)<br>[Midazolam / LC-MS/MS] |

**Figure 3.**
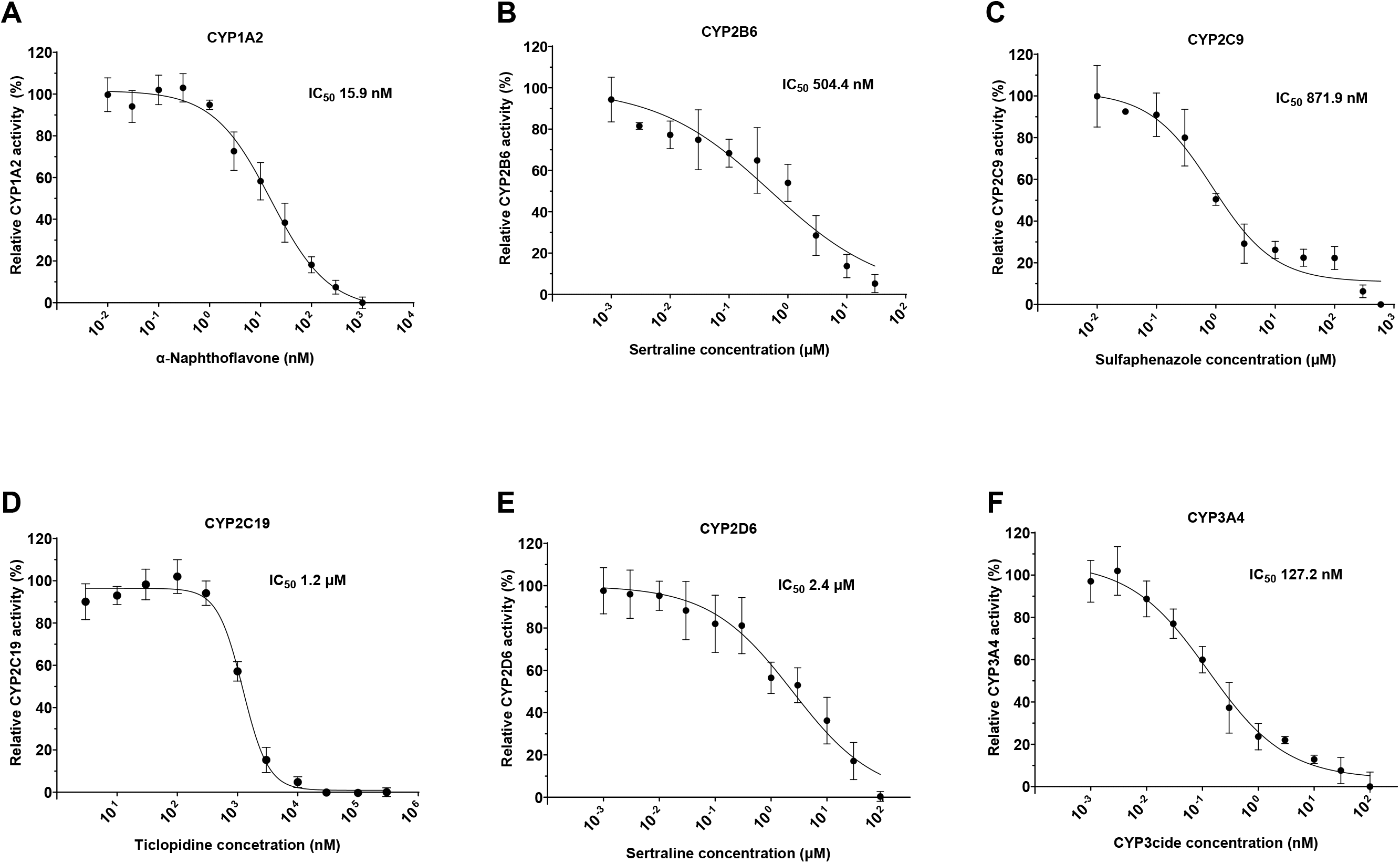
Dose dependent inhibition of CYPs using specific inhibitors. (A) CYP1A2 was inhibited by α-naphthoflavone after 10 minutes of pre-incubation. (B) CYP2B6 was inhibited by sertraline after 30 minutes of pre-incubation. (C) CYP2C9 was inhibited by sulfaphenazole after 10 minutes of pre-incubation. (D) CYP2C19 was inhibited by ticlopidine after 5 minutes of pre-incubation (E) CYP2D6 was inhibited by sertraline after 30 minutes of pre-incubation. (F) CYP3A4 was inhibited by CYP3cide after 5 minutes of pre-incubation. IC_50_values presented in the figure are for the biological replicate shown. All data have been collected as three biological replicates with average IC_50_values and standard deviations presented in Table S4.

### 4.4 Determination of the type of inhibition for select CYP-inhibitor pairs

After determining the IC_50_for each of the specific inhibitors, the utility of the developed method to determine the inhibition type for select CYP-inhibitor pairs was explored. The same concentrations of the substrate used in the previous Michaelis-Menten kinetics determinations were used again along with varying concentrations of the inhibitor. Fig. 4A and 4B show the resultant data for inhibition of CYP2C9 by sulfaphenazole and inhibition of CYP2C19 by ticlopidine, respectively. Sulfaphenazole inhibition of CYP2C9 yielded no change in the V_max_as the concentration of inhibitor changed suggesting competitive inhibition and comparisons using Akaike’s Information Criterion selected competitive inhibition as the model that most likely produced the data. The average CYP2C9 inhibition constant, K_*i*_, of sulfaphenazole was 2.06 ± 0.3μM with an average V_max_of 0.37 ± 0.0 nmol/min (Table S5 and S6). In the case of ticlopidine inhibition of CYP2C19, V_max_clearly decreases with increasing concentrations of ticlopidine suggesting non-competitive inhibition.

**Figure 4.**
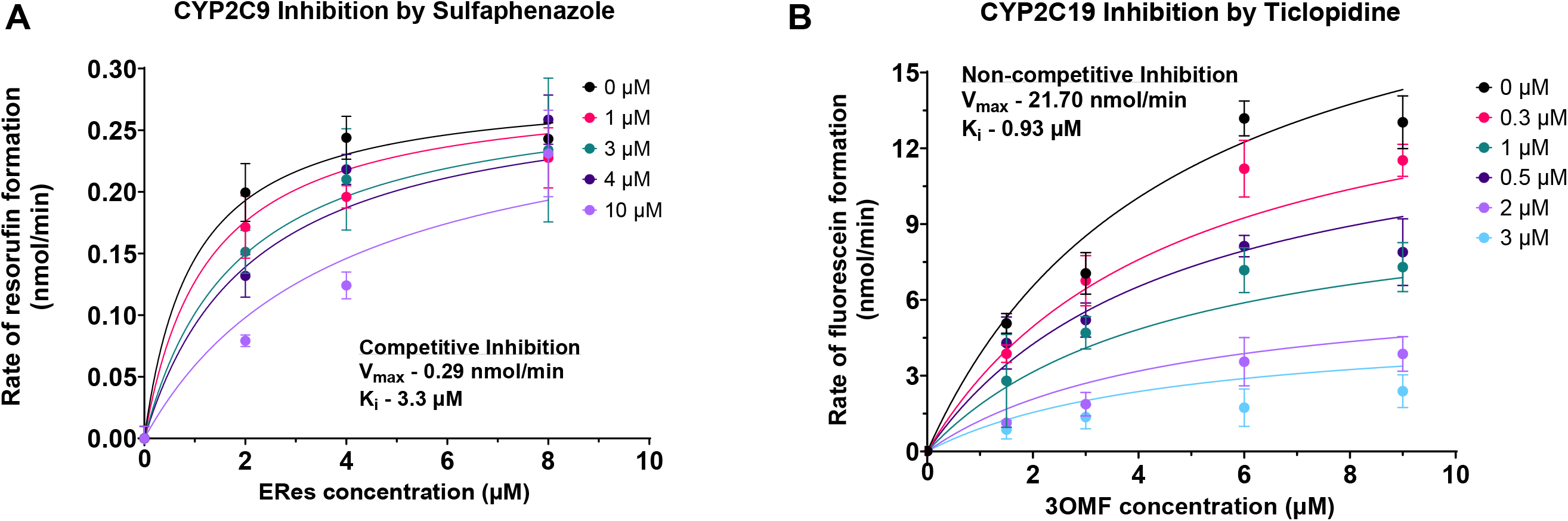
Type of inhibition for CYP2C19 with ticlopidine and CYP2C9 with sulfaphenazole. (A) CYP2C9 was inhibited by sulfaphenazole in a competitive way with V_max_0.29 nmol/min (95% CI – 0.26-0.31) and K_*i*_of 3.3 μM (95% CI – 1.7-6.0). (B) CYP2C19 was inhibited by ticlopidine in a non-competitive way with V_max_21.70 nmol/min (95% CI – 19.35-24.69) and K_i_of 0.93 μM (95% CI – 0.81-1.06). K_*i*_and V_max_values presented in the figure are for the biological replicate shown. All the experiments have been performed as three biological replicates with average and standard deviation of K_i_and V_max_presented in Table S5 and S6.

Indeed, comparisons using Akaike’s Information Criterion also determine that non-competitive inhibition is the model most likely to have generated the data and fits to mixed inhibition yield an alpha value close to one also suggestive of non-competitive inhibition. The average inhibition constant K_*i*_was found to be 1.05 ± 0.3μM with an average V_max_of 23.4 ± 5.5 nmol/min.

## 5 Discussion

In humans, CYPs play a critical role in xenobiotic metabolism and drug-drug interactions such that the determination of inhibition by and metabolism of potential therapeutics is an important step in the drug development pipeline. Therefore, a high-throughput, kinetic fluorogenic assay system that can be used for the determination of inhibition or activation of the CYPs most important for xenobiotic metabolism in humans is of critical importance (Leonard and Koffas, 2007, Nelson, 2009, Quehl et al., 2016).

Herein we report the development of pFluor50, a set of high-throughput fluorogenic CYP assays using the commercially available CYPExpress™ reagents which provide reproducible kinetic data suitable for screening compounds for CYP inhibition or activation, determining the time-dependence of that inhibition or activation, and the inhibition type. To our knowledge the fluorogenic method described herein is one of the few that has been demonstrated to determine mechanism-based or time dependent CYP inhibition and also type of inhibition. Each CYP enzyme and master-mix was optimized to include the appropriate specific reagents needed to provide linear responses within the time window selected when paired with the appropriate fluorogenic substrates (see Table 1 and supplementary materials). Additionally, the total incubation times and Ex:Em values were also optimized. Further, the parameters for the plate reader such as the speed with which reads were collected, the number of measurements for every read, the total time interval between two consecutive reads, the height at which the fluorescence was detected, and the fluorescence gain values were determined and then fixed to ensure reproducible readings every time the assay was run (Table S1) and to aid in method transfer so the assays can be easily replicated by others. All parameters were optimized to ensure the linearity of the increase in fluorescence due to product formation within the linear phase of the enzyme kinetics. Optimization was performed by empirical methods and the appropriate conditions are reported here and in the supplementary materials. The optimization of these plate reader parameters along with kinetic parameters makes the method easily reproducible and transferable and it can be used effectively for determining CYP inhibition/activation.

Because the *K*_M_ can depend on many factors including pH, temperature, and ionic strength (Hochachka and Lewis, 1971), the methods were developed at a constant pH value of 7.4 and a constant temperature of 37 °C (physiological pH and temperature) (Burton, 1935, Proksch, 2018). Although *K*_M_ is independent of enzyme concentration V_max_is dependent on enzyme concentration (Nelson et al., 2008). In this study we have used a commercially available proprietary enzyme mix (which contains the CYP enzymes along with the additional enzymes NAPDH oxidoreductase and G6PDH as well as MgCl_2_) wherein the supplier does not provide the information on the exact quantity of each component. Therefore, we have included the amount of the CYPExpress™ reagent used in each case rather than the concentration of each individual component. Nonetheless, the *K*_M_ values obtained using the developed assays in this study were quite similar to the reported *K*_M_ values for the CYP-substrate pairs when values were available in the literature (Table 1) as was the case for CYP1A2, CYP2B6, CYP2C19 and CYP3A4 (Appiah-Opong et al., 2007, Murayama et al., 2004, Stresser, 2004, Sudsakorn et al., 2007). To the best of our knowledge, this is the first report of the use of Eres as a substrate for CYP2C9 and CYP2D6. In all cases where the fluorescent product was commercially available, the V_max_was calculated as the rate of formation of the product (Table 1). However, fluorescein benzyl ester (the fluorescent product following the cleavage of dibenzyl fluorescein) is not commercially available, therefore the V_max_for CYP3A4 cleavage of dibenzyl fluorescein has been reported as the rate of change in relative fluorescence units (RFU) per minute.

The newly developed pFluor50 methods were further validated by determining the IC_50_for known inhibitors of each of the six CYPs. One of the major factors that was considered when performing the inhibition studies was the selection of the solvents used to dissolve the inhibitors. The solvent choice was found to be critical because some CYPs are very sensitive to some solvents, demonstrating a significant loss in activity even in the presence of 0.3-1% final solvent concentration (Busby et al., 1999, Li et al., 2010). Interestingly, it was also found that the inhibition potency of specific CYPs by the selected inhibitors (some of which are FDA approved drugs) was shown to be time dependent. Therefore, the pre-incubation time between the inhibitor and the CYP was first optimized prior to IC_50_determination. It was found that some inhibitors such as ticlopidine, CYP3cide and sertraline were time dependent while α-naphthoflavone and sulfaphenazole were time independent inhibitors (Fig. 2). Based on our analysis, the inhibition times for these inhibitors were optimized. CYP2B6 and CYP2D6 can metabolize sertraline with different potencies at low or high concentrations (Greenblatt et al., 1999, Kobayashi et al., 1999, Palacharla et al., 2018, Xu et al., 1999) and at short incubation times the data were highly variable (likely due to metabolite formation and inhibition by metabolites) so a 30 minute pre-incubation time was selected for CYP2B6 and CYP2D6 inhibition by sertraline. Similarly, incubation times were optimized for ticlopidine, CYP3cide, α-naphthoflavone and sulfaphenazole.

After optimizing the pre-incubation time, CYP inhibition assays were performed using the specific inhibitors. The IC_50_values determined in this study (Fig. 3) were largely similar to those reported in the literature (Sudsakorn et al., 2007, Thelingwani et al., 2012, Walsky et al., 2006, Khan et al., 2007, Sproule et al., 1997, Tseng et al., 2014). It should be noted that although the pFluor50 assay methods differed from the methods used in the literature (which were traditional LC-MS/MS methods) and the substrates were also different, the determined IC_50_values were still consistent. It is possible, if not likely, that the small differences that were observed could be due to differences in the pre-incubation times as these were often not noted in the literature and the most significant observed differences were in the values calculated for the inhibitors that were time dependent. Nonetheless, this demonstrates the robustness of the developed methods in comparison to the traditional methods and confirms that the developed assays are suitable for screening and characterization of drugs in the drug development pipeline. Finally, the type of inhibition was determined for CYP2C9 inhibition by sulfaphenazole and CYP2C19 inhibition by ticlopidine. This further demonstrates that the methods developed are appropriate for not only determining inhibition but also characterizing the type of inhibition.

Additionally, because the CYPExpress™ reagents were selected as the source of the enzyme, further studies of metabolite production and mechanism-based inhibition would be simplified. Low-speed centrifugation could be used to remove the enzymes for the metabolite production studies, facilitating sample preparation prior to mass spectrometric determination of the metabolites produced. The same procedure could also be used to retain the enzymes allowing one to wash away the inhibitors prior to activity determination using this method to expedite the confirmation of mechanism-based inhibition.

Overall, fluorogenic assays are advantageous in several aspects compared to other more traditional CYP metabolism assays such as LC-MS/MS or HPLC based methods. They tend to cost less, require less expensive and more readily available instrumentation, and require less expertise in instrument handling. Further, fluorogenic assays are easily adaptable to high-throughput methods making them useful for screening a large number of drug candidates at a time. The pFluor50 assay system has been shown to provide comparable results to previously reported methods while providing real-time kinetic reads that enable further characterization of the observed inhibition, such as the time-dependence of the inhibition and the inhibition type (Table 1-4; Table S1, S3-S6).

Here, pFluor50, a novel fluorogenic assay system is described that has been developed to determine the inhibition of six human CYPs using the commercially available CYPExpress™ system. In this assay system, we have used Eres as a novel fluorogenic substrate for CYP2C9 and CYP2D6. The pFluor50 system can be used to provide data critical to the early stages of the drug development process and is among the few fluorogenic methods that have been shown to determine the time dependence of inhibition which can provide insight into the mechanism of action of the inhibitors as well as determine the type of inhibition. The approach outlined herein can be used to provide data in a high-throughput manner (faster than HPLC etc.) on six different human CYPs (CYP1A2, CYP2B6, CYP2C9, CYP2C19, CYP2D6 and CYP3A4) that are responsible for the metabolism of >90% of xenobiotics. Known CYP-specific inhibitors were used to validate the methods and demonstrate the utility of the methods for determining type of inhibition. The developed methods can therefore be used for rapid screening of drug candidates for inhibition or activation of CYP metabolism and further characterize any observed.

## 6 Acknowledgments

None.

## 10 Authors contribution

Conceptualization -PS and CM Data curation – PS

Funding Acquisition (resources) – SE and CM

Investigation – PS, AR, SR, and AB

Methodology -PS

Data acquisition -PS, AR, and AB

Formal analysis -PS and CM

Writing -original draft preparation – PS

Writing -reviewing and editing – PS, CM, TL, SE

Supervision – CM and SE

Project administration – CM and SE

## 13 Supplementary material

The supplementary file with tables and figures has been attached separately as a single word file.

## 14 Data Availability Statement

The raw data supporting the conclusions of this article will be made available by the authors upon request.

## References

appiah-opong, R., commandeur, J. N., van vugt-lussenburg, B. & vermeulen, N. P. 2007. Inhibition of human recombinant cytochrome P450s by curcumin and curcumin decomposition products. Toxicology, 235, 83–91.

atkins, W. M. 2005. Non-Michaelis-Menten kinetics in cytochrome P450-catalyzed reactions. Annu Rev Pharmacol Toxicol, 45, 291–310.

attar, M., shen, J., ling, K. H. & tang-liu, D. 2005. Ophthalmic drug delivery considerations at the cellular level: drug-metabolising enzymes and transporters. Expert Opin Drug Deliv, 2, 891–908.

ayrton, J., plumb, R., leavens, W. J., mallett, D., dickins, M. & dear, G. J. 1998. Application of a generic fast gradient liquid chromatography tandem mass spectrometry method for the analysis of cytochrome P450 probe substrates. Rapid Commun Mass Spectrom, 12, 217–24.

beam, A. & motsinger-reif, A. 2014. Beyond IC(50)s: Towards Robust Statistical Methods for in vitro Association Studies. J Pharmacogenomics Pharmacoproteomics, 5, 1000121.

bell, L., bickford, S., nguyen, P. H., wang, J., he, T., zhang, B., friche, Y., zimmerlin, A., urban, L. & bojanic, D. 2008. Evaluation of fluorescence- and mass spectrometry-based CYP inhibition assays for use in drug discovery. J Biomol Screen, 13, 343–53.

burton, A. C. 1935. Human calorimetry: II. The average temperature of the tissues of the body: three figures. The Journal of Nutrition, 9, 261–280.

busby, W. F., ackermann, J. M. & crespi, C. L. 1999. Effect of methanol, ethanol, dimethyl sulfoxide, and acetonitrile on in vitro activities of cDNA-expressed human cytochromes P-450. Drug Metab Dispos, 27, 246–9.

cohen, L. H., remley, M. J., raunig, D. & vaz, A. D. 2003. In vitro drug interactions of cytochrome p450: an evaluation of fluorogenic to conventional substrates. Drug Metab Dispos, 31, 1005–15.

deodhar, M., al rihani, S. B., arwood, M. J., darakjian, L., dow, P., turgeon, J. & michaud, V. 2020. Mechanisms of CYP450 Inhibition: Understanding Drug-Drug Interactions Due to Mechanism-Based Inhibition in Clinical Practice. Pharmaceutics, 12.

donahue, S. R., flockhart, D. A., abernethy, D. R. & ko, J. W. 1997. Ticlopidine inhibition of phenytoin metabolism mediated by potent inhibition of CYP2C19. Clin Pharmacol Ther, 62, 572–7.

donato, M. T., jimÉnez, N., castell, J. V. & gÓmez-lechÓn, M. J. 2004. Fluorescence-based assays for screening nine cytochrome P450 (P450) activities in intact cells expressing individual human P450 enzymes. Drug Metab Dispos, 32, 699–706.

galetin, A., gertz, M. & houston, J. B. 2010. Contribution of intestinal cytochrome p450mediated metabolism to drug-drug inhibition and induction interactions. Drug Metab Pharmacokinet, 25, 28–47.

greenblatt, D. J., von moltke, L. L., harmatz, J. S. & shader, R. I. 1999. Human cytochromes mediating sertraline biotransformation: seeking attribution. J Clin Psychopharmacol, 19, 489–93.

ha-duong, N. T., marques-soares, C., dijols, S., sari, M. A., dansette, P. M. & mansuy, D. 2001. Interaction of new sulfaphenazole derivatives with human liver cytochrome p450 2Cs: structural determinants required for selective recognition by CYP 2C9 and for inhibition of human CYP 2Cs. Arch Biochem Biophys, 394, 189–200.

hasler, J. A., estabrook, R., murray, M., pikuleva, I., waterman, M., capdevila, J., holla, V., helvig, C., falck, J. R. & farrell, G. 1999. Human cytochromes P450. Molecular aspects of medicine, 20, 1–137.

hinkson, I. V., madej, B. & stahlberg, E. A. 2020. Accelerating Therapeutics for Opportunities in Medicine: A Paradigm Shift in Drug Discovery. Front Pharmacol, 11, 770.

hochachka, P. W. & lewis, J. K. 1971. Interacting effects of pH and temperature on the K m values for fish tissue lactate dehydrogenases. Comp Biochem Physiol B, 39, 925–33.

jin, X., potter, B., luong, T. L., nelson, J., vuong, C., potter, C., xie, L., zhang, J., zhang, P., sousa, J., li, Q., pybus, B. S., kreishman-deitrick, M., hickman, M., smith, P. L., paris, R., reichard, G. & marcsisin, S. R. 2016. Pre-clinical evaluation of CYP 2D6 dependent drug-drug interactions between primaquine and SSRI/SNRI antidepressants. Malar J, 15, 280.

kabir, M. L., wang, F. & clayton, A. H. A. 2022. Intrinsically Fluorescent Anti-Cancer Drugs. Biology (Basel), 11.

khan, M., mohan, I. K., kutala, V. K., kumbala, D. & kuppusamy, P. 2007. Cardioprotection by sulfaphenazole, a cytochrome p450 inhibitor: mitigation of ischemiareperfusion injury by scavenging of reactive oxygen species. Journal of Pharmacology and Experimental Therapeutics, 323, 813–821.

kobayashi, K., ishizuka, T., shimada, N., yoshimura, Y., kamijima, K. & chiba, K. 1999. Sertraline N-demethylation is catalyzed by multiple isoforms of human cytochrome P-450 in vitro. Drug Metab Dispos, 27, 763–6.

laine, K., tybring, G., hÄrtter, S., andersson, K., svensson, J. O., widÉn, J. & bertilsson, L. 2001. Inhibition of cytochrome P4502D6 activity with paroxetine normalizes the ultrarapid metabolizer phenotype as measured by nortriptyline pharmacokinetics and the debrisoquin test. Clin Pharmacol Ther, 70, 327–35.

lamb, D. C., lei, L., warrilow, A. G., lepesheva, G. I., mullins, J. G., waterman, M. R. & kelly, S. L. 2009. The first virally encoded cytochrome p450. J Virol, 83, 8266–9.

le, S. B., holmuhamedov, E. L., narayanan, V. L., sausville, E. A. & kaufmann, S. H. 2006. Adaphostin and other anticancer drugs quench the fluorescence of mitochondrial potential probes. Cell Death Differ, 13, 151–9.

leonard, E. & koffas, M. A. 2007. Engineering of artificial plant cytochrome P450 enzymes for synthesis of isoflavones by Escherichia coli. Appl Environ Microbiol, 73, 7246–51.

li, D., han, Y., meng, X., sun, X., yu, Q., li, Y., wan, L., huo, Y. & guo, C. 2010. Effect of regular organic solvents on cytochrome P450-mediated metabolic activities in rat liver microsomes. Drug Metab Dispos, 38, 1922–5.

mckinnon, R. A., sorich, M. J. & ward, M. B. 2008. Cytochrome P450 part 1: multiplicity and function. Journal of pharmacy practice and research, 38, 55–57.

miller, V. P., stresser, D. M., blanchard, A. P., turner, S. & crespi, C. L. 2000. Fluorometric high-throughput screening for inhibitors of cytochrome P450. Ann N Y Acad Sci, 919, 26–32.

murayama, N., soyama, A., saito, Y., nakajima, Y., komamura, K., ueno, K., kamakura, S., kitakaze, M., kimura, H., goto, Y., saitoh, O., katoh, M., ohnuma, T., kawai, M., sugai, K., ohtsuki, T., suzuki, C., minami, N., ozawa, S. & sawada, J. 2004. Six novel nonsynonymous CYP1A2 gene polymorphisms: catalytic activities of the naturally occurring variant enzymes. J Pharmacol Exp Ther, 308, 300–6.

nayadu, S., behera, D., sharma, M., kaur, G. & gudi, G. 2013. Fluorescent probe based CYP inhibition assay: A high throughput tool for early drug discovery screening. Int Pharm Pharm Sci, 5, 303–307.

nelson, D. L., lehninger, A. L. & cox, M. M. 2008. Lehninger principles of biochemistry, Macmillan. NELSON, D. R.2009. The cytochrome p450 homepage. Hum Genomics, 4, 59–65.

nishiya, Y., hagihara, K., kurihara, A., okudaira, N., farid, N. A., okazaki, O. & ikeda, T. 2009. Comparison of mechanism-based inhibition of human cytochrome P450 2C19 by ticlopidine, clopidogrel, and prasugrel. Xenobiotica, 39, 836–43.

obach, R. S., cox, L. M. & tremaine, L. M. 2005. Sertraline is metabolized by multiple cytochrome P450 enzymes, monoamine oxidases, and glucuronyl transferases in human: an in vitro study. Drug Metab Dispos, 33, 262–70.

ogu, C. C. & maxa, J. L. 2000. Drug interactions due to cytochrome P450. Proc (Bayl Univ Med Cent), 13, 421–3.

okudaira, T., kotegawa, T., imai, H., tsutsumi, K., nakano, S. & ohashi, K. 2007. Effect of the treatment period with erythromycin on cytochrome P450 3A activity in humans. The journal of clinical pharmacology, 47, 871–876.

ortiz de montellano, P. R. 2013. Cytochrome P450-activated prodrugs. Future Med Chem, 5, 213–28.

otton, S. V., ball, S. E., cheung, S. W., inaba, T., rudolph, R. L. & sellers, E. M. 1996. Venlafaxine oxidation in vitro is catalysed by CYP2D6. Br J Clin Pharmacol, 41, 149–56.

palacharla, R. C., nirogi, R., uthukam, V., manoharan, A., ponnamaneni, R. K. & kalaikadhiban, I. 2018. Quantitative in vitro phenotyping and prediction of drug interaction potential of CYP2B6 substrates as victims. Xenobiotica, 48, 663–675.

paradise, E., chaturvedi, P. & ter-ovanesyan, E. 2007. Cytochrome P450 inhibition assays using traditional and fluorescent substrates. Curr Protoc Pharmacol, Chapter 7, Unit7.11.

park, E. J., park, K., durai, P., kim, K. Y., park, S. Y., kwon, J., lee, H. J., pan, C. H. & liu, K. H. 2022. Potent and Selective Inhibition of CYP1A2 Enzyme by Obtusifolin and Its Chemopreventive Effects. Pharmaceutics, 14.

proksch, E. 2018. pH in nature, humans and skin. J Dermatol, 45, 1044–1052.

quehl, P., hollender, J., schÜÜrmann, J., brossette, T., maas, R. & jose, J. 2016. Co-expression of active human cytochrome P450 1A2 and cytochrome P450 reductase on the cell surface of Escherichia coli. Microbial Cell Factories, 15, 1–15.

raunio, H., pentikainen, O. & juvonen, R. O. 2020. Coumarin-Based Profluorescent and Fluorescent Substrates for Determining Xenobiotic-Metabolizing Enzyme Activities In Vitro. Int J Mol Sci, 21.

sadler, N. C., nandhikonda, P., webb-robertson, B. J., ansong, C., anderson, L. N., smith, J. N., corley, R. A. & wright, A. T. 2016. Hepatic Cytochrome P450 Activity, Abundance, and Expression Throughout Human Development. Drug Metab Dispos, 44, 984–91.

sproule, B. A., otton, S. V., cheung, S. W., zhong, X. H., romach, M. K. & sellers, E. M. 1997. CYP2D6 inhibition in patients treated with sertraline. J Clin Psychopharmacol, 17, 102–6.

stresser, D. M. 2004. High-Throughput Screening of Human Cytochrome P450 Inhibitors Using Fluorometric Substrates: Methodology for 25 Enzyme/Substrate Pairs. Optimization in drug discovery: in vitro methods, 215–230.

stresser, D. M., blanchard, A. P., turner, S. D., erve, J. C., dandeneau, A. A., miller, V. P. & crespi, C. L. 2000. Substrate-dependent modulation of CYP3A4 catalytic activity: analysis of 27 test compounds with four fluorometric substrates. Drug Metab Dispos, 28, 1440–8.

sudsakorn, S., skell, J., williams, D. A., o’shea, T. J. & liu, H. 2007. Evaluation of 3-O-methylfluorescein as a selective fluorometric substrate for CYP2C19 in human liver microsomes. Drug Metab Dispos, 35, 841–7.

sukumaran, S. M., potsaid, B., lee, M. Y., clark, D. S. & dordick, J. S. 2009. Development of a fluorescence-based, ultra high-throughput screening platform for nanoliter-scale cytochrome p450 microarrays. J Biomol Screen, 14, 668–78.

sun, D., gao, W., hu, H. & zhou, S. 2022. Why 90% of clinical drug development fails and how to improve it? Acta Pharm Sin B, 12, 3049–3062.

thelingwani, R. S., dhansay, K., smith, P., chibale, K. & masimirembwa, C. M. 2012. Potent inhibition of CYP1A2 by Frutinone A, an active ingredient of the broad spectrum antimicrobial herbal extract from P. fruticosa. Xenobiotica, 42, 989–1000.

thomford, N. E., dzobo, K., chopera, D., wonkam, A., maroyi, A., blackhurst, D. & dandara, C. 2016. In Vitro Reversible and Time-Dependent CYP450 Inhibition Profiles of Medicinal Herbal Plant Extracts Newbouldia laevis and Cassia abbreviata: Implications for Herb-Drug Interactions. Molecules, 21.

tseng, E., walsky, R. L., luzietti, R. A., harris, J. J., kosa, R. E., goosen, T. C., zientek, M. A. & obach, R. S. 2014. Relative contributions of cytochrome CYP3A4 versus CYP3A5 for CYP3A-cleared drugs assessed in vitro using a CYP3A4-selective inactivator (CYP3cide). Drug Metab Dispos, 42, 1163–73.

ung, Y. T., ong, C. E. & pan, Y. 2018. Current High-Throughput Approaches of Screening Modulatory Effects of Xenobiotics on Cytochrome P450 (CYP) Enzymes. High Throughput, 7.

walsky, R. L., astuccio, A. V. & obach, R. S. 2006. Evaluation of 227 drugs for in vitro inhibition of cytochrome P450 2B6. J Clin Pharmacol, 46, 1426–38.

walsky, R. L., obach, R. S., hyland, R., kang, P., zhou, S., west, M., geoghegan, K. F., helal, C. J., walker, G. S., goosen, T. C. & zientek, M. A. 2012. Selective mechanism-based inactivation of CYP3A4 by CYP3cide (PF-04981517) and its utility as an in vitro tool for delineating the relative roles of CYP3A4 versus CYP3A5 in the metabolism of drugs. Drug Metab Dispos, 40, 1686–97.

xu, Z. H., wang, W., zhao, X. J., huang, S. L., zhu, B., he, N., shu, Y., liu, Z. Q. & zhou, H. H. 1999. Evidence for involvement of polymorphic CYP2C19 and 2C9 in the N-demethylation of sertraline in human liver microsomes. Br J Clin Pharmacol, 48, 416–23.

yan, Z. & caldwell, G. W. 2021. Cytochrome P450: In Vitro Methods and Protocols, Springer.

zanger, U. M. & schwab, M. 2013. Cytochrome P450 enzymes in drug metabolism: regulation of gene expression, enzyme activities, and impact of genetic variation. Pharmacol Ther, 138, 103–41.

zhao, M., ma, J., li, M., zhang, Y., jiang, B., zhao, X., huai, C., shen, L., zhang, N., he, L. & qin, S. 2021. Cytochrome P450 Enzymes and Drug Metabolism in Humans. Int J Mol Sci, 22.

