## Supplemental Material for "Development and characterization of pFluor50, a fluorogenic-based kinetic assay system for high-throughput inhibition screening and characterization of time-dependent inhibition and inhibition type for six human CYPs"

### *Supplementary Material*

#### 1 Supplementary Figures and Tables

##### 1.1 Supplementary Tables

**Supplementary Table 1. Synergy H1 Biotek plate parameters used during fluorogenic CYP assays**

| CYP-substrate | Speed | Delay (msec) | Measurements per read | Time interval (min) | Height (mM) | Gain | Reading time for linear region (min) |
|---|---|---|---|---|---|---|---|
| CYP1A2 | Normal | 100 | 45/read | 1.5 | 9.75 | 100 | 0-10 |
| CYP2B6 | Normal | 100 | 75/read | 1.5 | 9.75 | 130 | 0-60 |
| CYP2C9 | Normal | 100 | 100/read | 2 | 9.75 | 100 | 0-40 |
| CYP2C19 | Normal | 100 | 20/read | 0.5 | 9.5 | 68 | 0-20 |
| CYP2D6 | Normal | 100 | 75/read | 1.5 | 9.75 | 100 | 0-75 |
| CYP3A4 | Normal | 100 | 17/read | 1 | 9.5 | 100 | 15-35 |

**Supplementary Table 2. Michaelis Menten kinetic parameters for different CYP-substrate pairs**

| CYP-Substrate | $K_M$ – repeat 1 (95% CI) | $K_M$ – repeat 2 (95% CI) | $K_M$ – repeat 3 (95% CI) | Average $K_M$ |
|---|---|---|---|---|
| CYP1A2-Eres | 0.51 µM (0.41-0.64) | 0.52 µM (0.41-0.66) | 1.3 µM (1.0-1.6) | 0.78±0.3 µM |
| CYP2B6-Bzres | 23.4 µM (19.3-27.0) | 28.6 µM (23.5-35.1) | 18.9 µM (12.8-29.7) | 23.6±4.8 µM |
| CYP2C9-Eres | 0.49 µM (0.43-0.56) | 0.49 µM (0.36-0.65) | 0.28 µM (0.24-0.33) | 0.42±0.1 µM |
| CYP2C19-3OMF | 2.8 µM (2.4-3.2) | 4.1 µM (3.4-4.9) | 2.4 µM (1.5-3.9) | 3.1±0.8 µM |
| CYP2D6-Eres | 3.6 µM (3.0-4.3) | 3.1 µM (2.2-4.3) | 2.0 µM (1.4-2.9) | 2.91±0.8 µM |
| CYP3A4-DBF | 2.0 µM (1.7-2.3) | 1.3 µM (1.0-1.5) | 0.77 µM (0.60-0.99) | 1.35±0.6 µM |

**Supplementary Table 3. $V_{max}$ (rate of product formation) for different CYP-substrate pairs**

| CYP-Substrate | $V_{max}$ – repeat 1 (95% CI) | $V_{max}$ – repeat 2 (95% CI) | $V_{max}$ – repeat 3 (95% CI) | Average $V_{max}$ |
|---|---|---|---|---|
| CYP1A2-Eres | 1.86 nmol/min (1.75-1.99) | 1.45 nmol/min (1.34-1.58) | 2.2 nmol/min (1.99-2.47) nm | 1.83 ± 0.4 nmol/min |
| CYP2B6-Bzres | 436.6 pmol/min (410.7-466.2) | 573.6 pmol/min (513.1-648.9) | 456.7 pmol/min (368.7-607.1) | 620.2 ± 210.8 pmol/min |
| CYP2C9-Eres | 0.42 nmol/min (0.40-0.43) | 0.41 nmol/min (0.38-0.44) | 0.44 nmol/min (0.42-0.45) | 0.42 ± 0.0 nmol/min |
| CYP2C19-3OMF | 18.9 nmol/min (18.0-19.8) µM | 21.5 nmol/min (20.1-23.1) µM | 17.7 nmol/min (14.8-22.7) µM | 19.3 ± 1.9 nmol/min |
| CYP2D6-Eres | 1.11 nmol/min (1.06-1.17) | 0.80 µM nmol/min (0.73-0.86) | 1.11 µM nmol/min (0.99-1.12) | 1.00 ± 0.1 nmol/min |
| CYP3A4-DBF | 657.9 (RFU/time) (622.9-697.4) | 569.2 (RFU/time) (536.1-606.7) | 905.0 (RFU/time) (857.9-955.0) | 710.7 ± 174.0 (RFU/time) |

**Supplementary Table 4. Average $IC_{50}$ values for each CYP-inhibitor pair (three biological replicates)**

| CYP-inhibitor | $IC_{50}$ – repeat 1 (95% CI) | $IC_{50}$ – repeat 2 (95% CI) | $IC_{50}$ – repeat 3 (95% CI) | Average $IC_{50}$ (µM) |
|---|---|---|---|---|
| CYP1A2- α-Napthaflavone | 0.51 µM (0.41-0.64) µM | 0.82 µM (0.62-1.0) µM | 1.1 µM (0.81-1.4) µM | 0.81 ± 0.3 |
| CYP2B6-Sertraline | 0.367 µM (19.3-27.0) µM | 0.504 µM (32.1-46.8) µM | 0.577 µM (20.7-30.2) µM | 0.48 ± 0.1 |
| CYP2C9-Sulfaphenazole | 0.49 µM (0.43-0.56) µM | 0.28 µM (0.25-0.30) µM | 0.72 µM (0.59-0.86) µM | 0.49 ± 0.2 |
| CYP2C19-Ticlopidine | 2.8 µM (2.4-3.2) µM | 4.1 µM (3.4-4.9) µM | 2.4 µM (1.5-3.9) µM | 3.1 ± 0.8 |
| CYP2D6-Sertraline | 2.4 µM (1.6-3.5) µM | 1.8 µM (1.1-2.9) µM | 2.0 µM (1.0-3.6) µM | 2.06 ± 0.3 |
| CYP3A4-CYP3cide | 2.0 µM (1.7-2.3) µM | 1.3 µM (1.0-1.5) µM | 0.77 µM (0.60-0.99) µM | 1.35 ± 0.6 |

**Supplementary Table 5. Average $K_i$ for type of inhibition analysis (three biological replicates)**

| CYP-inhibitor | $K_i$ – repeat 1 (95% CI) | $K_i$ – repeat 2 (95% CI) | $K_i$ – repeat 3 (95% CI) | Average $K_i$ (µM) |
|---|---|---|---|---|
| CYP2C9-Sulfaphenazole | 3.3 µM (1.7-6.0) | 5.2 µM (3.6-6.1) | 5.1 µM (3.9-6.1) | 2.06 ± 0.3 |
| CYP2C19-Ticlopidine | 0.93 µM (0.80-1.06) | 1.40 µM (0.86-3.3) | 0.82 µM (0.62-1.1) | 1.05 ± 0.3 |

**Supplementary Table 6. Average $V_{max}$ for type of inhibition analysis (three biological replicates)**

| CYP-inhibitor | $V_{max}$ – repeat 1 (95% CI) | $V_{max}$ – repeat 2 (95% CI) | $V_{max}$ – repeat 3 (95% CI) | Average $V_{max}$ |
|---|---|---|---|---|
| CYP2C9-Sulfaphenazole | 0.29 nmol/min (1.6-3.5) | 0.4 nmol/min (0.34.1-0.51) | 0.43 µM nmol/min (0.36-0.51) | 0.37 ± 0.0 nmol/min |
| CYP2C19-Ticlopidine | 21.7 nmol/min (19.35-24.69) | 29.6 nmol/min (27.9-31.5) | 18.9 nmol/min (14.6-26.5) | 23.4 ± 5.5 nmol/min |

#### 1.2 Supplementary Figures

**Supplementary Figure 1. Plate layout for Michaelis-Menten analysis using $K_M$ substrate**

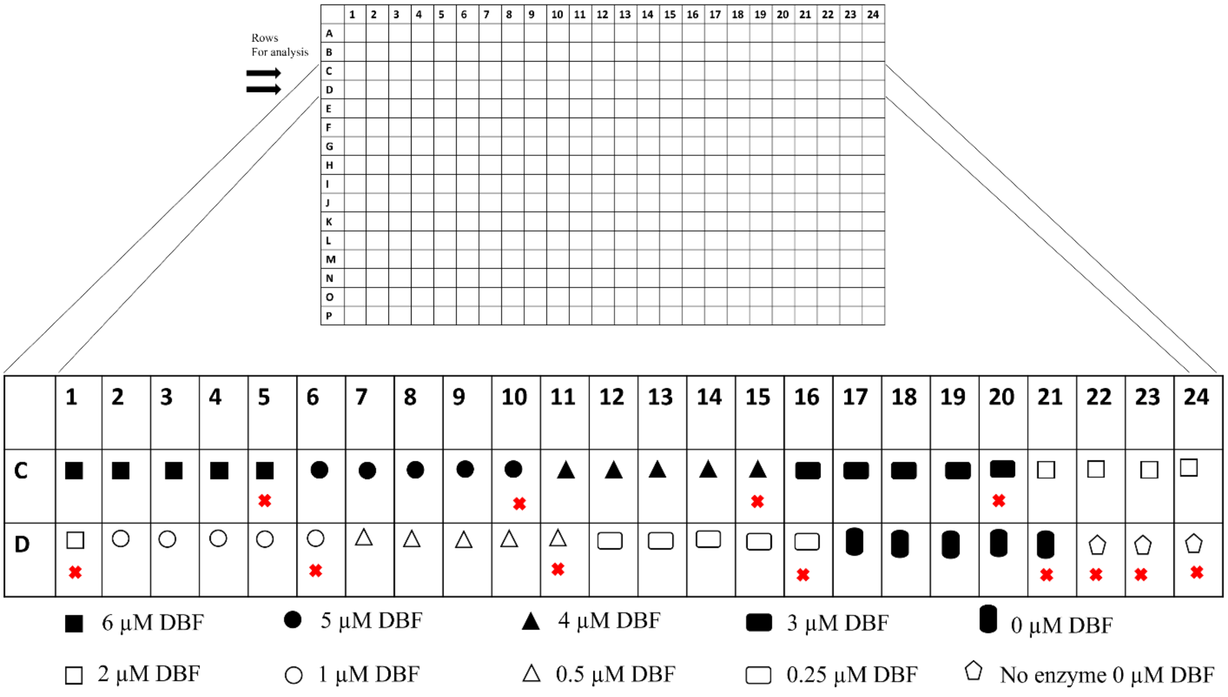

384 blackwell opti-plate layout for performing Michaelis-Menten analysis using CYP3A4 enzyme and DBF substrate. Different concentrations of substrate DBF were used in 4 replicates and a blank was used without the enzyme (✖) but with the substrate for each concentration of DBF. (⬠) Another control was used in which no enzyme as well as no inhibitor was used. This was further used to normalize all the readings.

**Supplementary Figure 2. CYP-substrate pairs used in fluorogenic assay development**

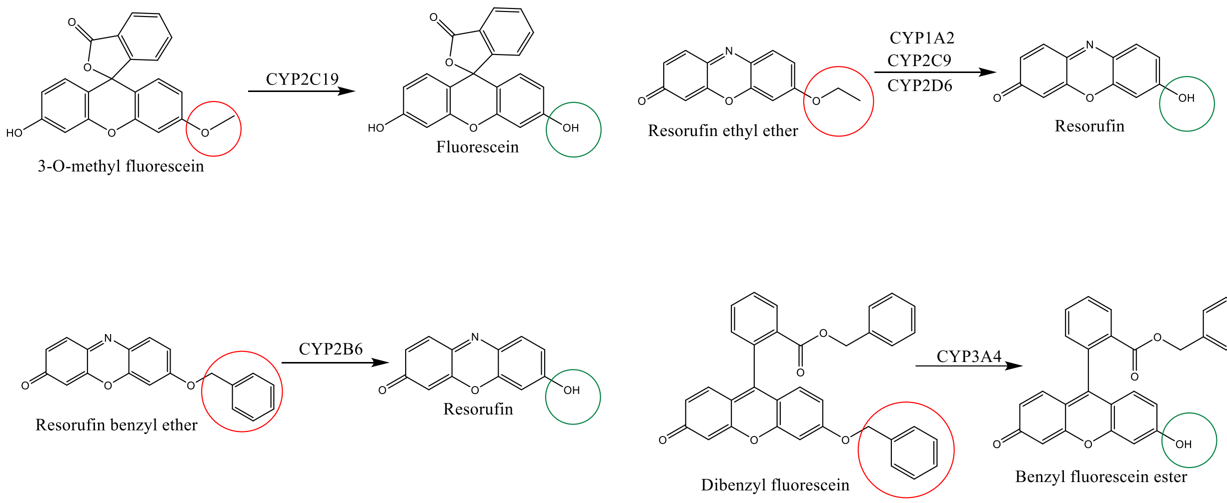


Structures of substrates and their final products used in fluorogenic assay development. The groups circled in red are metabolized to the ones in green by the specific CYP450 enzymes.

**Supplementary Figure 3. Calibration curves for Resorufin and Fluorescein**

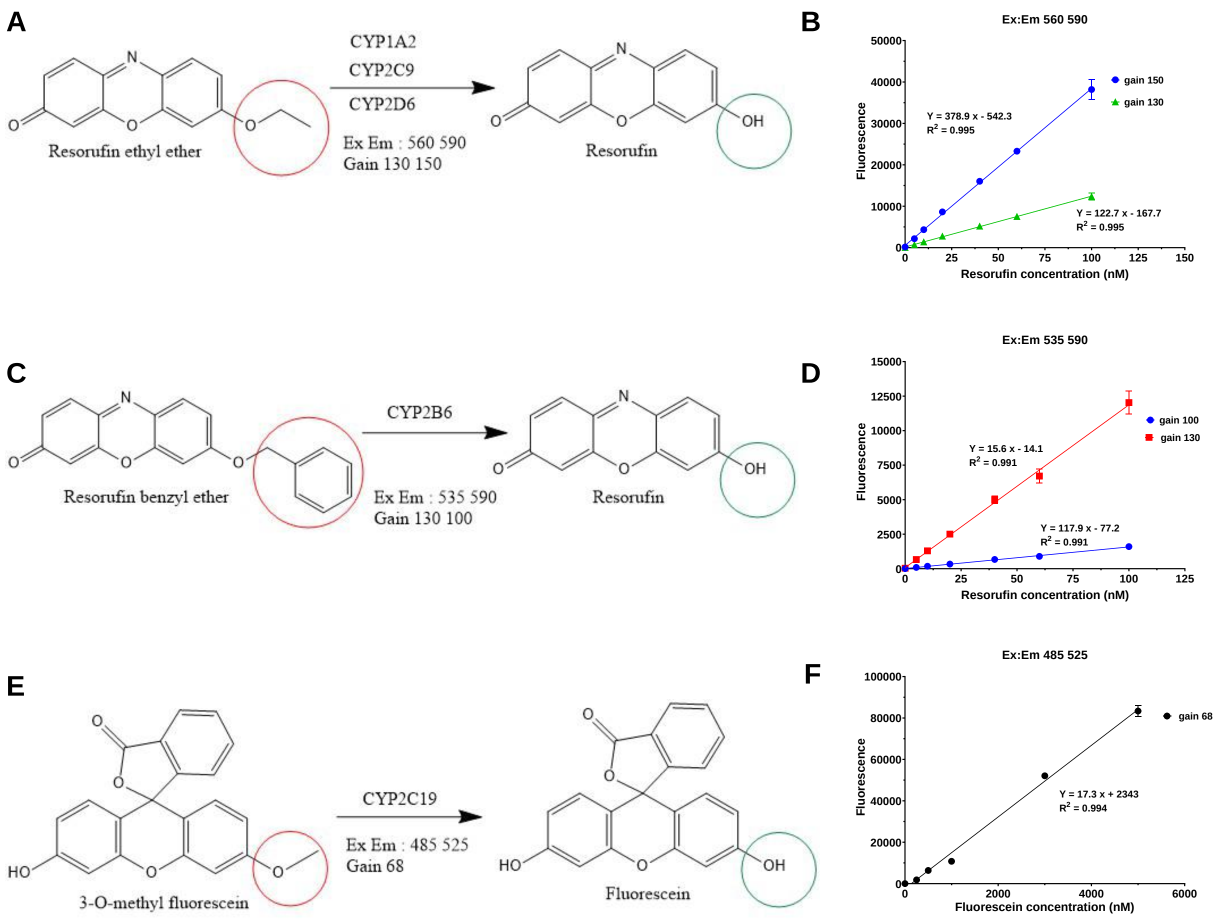


(A) Ex:Em (with gain values 150 and 130) values for formation of resorufin from resorufin ethyl ester by CYP1A2, CYP2D6 and CYP2C9. (B) Fluorescence values for resorufin (0 – 100 nM) at Ex:Em 560:590 (gain 150 and 130). (C) Ex:Em (with gain values 100 and 130) values for formation of formation of resorufin from resorufin benzyl ester by CYP2B6. (D) Fluorescence calibration curve for resorufin (0 – 100 nM) at Ex:Em 535:590 (gain 100 and 130). (E) Ex:Em (with gain value 68) values for formation of fluorescein from 3-O-Methyl fluorescein by CYP2C19. (F) Fluorescence calibration curve for fluorescein ( 0 – 5000 nM) at Ex:Em 485:520 (gain 68).
